# Maltose attenuates the virulence of *Salmonella* Typhimurium by impairing its entry into epithelial cells

**DOI:** 10.1101/2025.05.06.652415

**Authors:** Kirti Parmar, S Jayashree, Dipshikha Chakravortty

**Affiliations:** Department of Microbiology and Cell Biology, Division of Biological Sciences, Indian Institute of Science, Bangalore, India; Adjunct Faculty, School of Biology, Indian Institute of Science Education and Research, Thiruvananthapuram, India

**Keywords:** *Salmonella*, maltose, intestinal epithelial cells, adhesion, fimbriae

## Abstract

The composition of nutrients in the intestine defines a niche for colonising gut pathogens. The lack of nutrients suitable for pathogens due to competition from resident intestinal microbiota or dietary preferences leads to colonisation failure by pathogens. In this study, we investigated the impact of the disaccharide sugar maltose on *Salmonella* Typhimurium (STM) pathogenicity during the early phases of infection in C57BL/6 mice and human colon carcinoma cells (Caco-2). We found that supplementation of maltose at lower concentrations inhibited STM colonisation in the ileum of mice. To understand this, we investigated the role of maltose metabolism in human colon epithelial (Caco-2) cells. Deleting the maltose metabolism gene (malQ) increased adhesion to Caco-2 cells. The increased adhesion was due to increased expression of type 1 fimbriae. Inhibiting the type 1 fimbriae-mediated binding to host epithelial cells by incubating with mannose resulted in similar adhesion of STM WT and STM Δ*malQ.* We further identified that *malQ* regulates the adhesion of Salmonella through the sigma factor RpoE. Overall, *malQ* in *Salmonella* inhibits infection in epithelial cells by reducing its adhesion to host epithelial cells.

## Introduction

Foodborne pathogens encounter various stresses inside and outside the host (1). Pathogen growth often occurs in an environment with limited nutrients inside the host, owing to competition from resident intestinal microbiota in the host intestine (2). Bacteria, such as *Salmonella,* can successfully modulate their metabolism in response to the availability of nutrients. *Salmonella enterica* serovar Typhimurium causes self-limiting gastroenteritis in humans and causes bacteraemia in immunocompromised individuals (3).

Successful *Salmonella* colonisation depends on utilising multiple carbon sources, including glycerol, fatty acids, N-acetylglucosamine, gluconate, glucose, lactate, and arginine (4). Maltose in the intestine plays a pivotal role in the growth and pathogenesis of various bacteria, both beneficial and detrimental. Maltose is a commonly present carbohydrate source in food products like sweet potatoes and barley, and its concentration increases during cooking and fermentation (5). Compared with glucose, sugars such as maltose/maltodextrins are preferred for growth on poor nitrogen sources (6). The utilisation of maltose is also crucial for colonisation of the mouse intestine by *E.coli* (*7*).

In contrast, maltose derivatives reduce the biofilm production and adhesion of *Pseudomonas aeruginosa* non-swarming mutants (8). In zebrafish and crucian carp, maltose modulates immunity against *Vibrio alginolyticus* and *Aeromonas sobrial,* respectively, by increasing lysozyme activity and expression (9, 10). Lastly, in *Vibrio cholerae,* maltose inhibits cholera toxin secretion and reduces pathogenicity in newborn mice (11). Maltodextrins promote *Salmonella* survival in the bone marrow-derived macrophages by reducing NADPH expression and recruitment to the *Salmonella* containing vacuole. *In vivo,* maltodextrins degrade the intestinal mucosa of mice, increasing the cecal load (12). These observations highlight the diverse roles of maltose and maltodextrin in bacterial growth and pathogenesis. However, it remains unclear how maltose, a carbohydrate, affects *Salmonella’s* virulence factors. A strain deficient in the maltose hydrolysis gene (*malQ*) was constructed further to study the link between maltose metabolism and virulence. This study aimed to understand the modulation of *Salmonella* virulence with respect to maltose metabolism.

## Materials and Methods

### Bacteria and growth conditions

Wild-type *Salmonella* Typhimurium 14028s (STM WT) was obtained from Professor Michael Hensel (Universität Osnabrück). STM WT was revived from glycerol stock kept at -80°C and streaked on LB-agar; the respective antibiotic-containing plates were used for the knockout and complement strains. All the bacterial strains were grown in LB supplemented with the required antibiotics at 37°C and 170 rpm overnight. The strain containing pkD46 was grown at 30°C and 170 rpm. For induction of the complement strain, 0.2% (13.3 mM) L-arabinose was used during the bacterial subculture. A list of bacterial strains used in the study is attached in Supplementary Table 1.

### Construction of knockout strains

The gene knockout strains were prepared using a one-step chromosomal gene inactivation protocol developed by Datsenko and Wanner (13). In this protocol, knockout primers were designed to be 40 bp homologous to the gene to be knocked out, with a 20 bp extension for the antibiotic cassette. The gene coding for the antibiotic is amplified from the pKD3 or pKD4 plasmid, and through homologous recombination, the gene is replaced by an antibiotic cassette using electroporation. Amplified cassettes were inserted into STM pKD46, which has a lambda red recombinase system under an arabinose-inducible promoter. For electroporation, STM pKD46 bacterial cells were grown to the log phase, and the absorbance was adjusted to 0.3. The cells were then made electrocompetent by washing with MilliQ water and 10% glycerol at 4°C. The knockout strains were screened by plating the bacteria on an antibiotic-containing plate and performing PCR on the obtained colonies. The primers used in the study are attached in Supplementary Table 2.

### Construction of complement strains

*malQ* gene amplified from *Salmonella* Typhimurium and the pBAD vector was digested with KpnI and PstI at 37°C for 1 hour. Digested products were gel purified and ligated with T4 DNA ligase (NEB). Ligated products were transformed into *E.coli* Tg1 competent cells using the heat shock method. Transformed cells were plated on the LB-chloramphenicol and incubated overnight at 37°C. Plasmid isolation was performed from transformants. Cloning was confirmed by double digestion. After successful confirmation, the clone was transformed into STM Δ*malQ*.

Construction of reporter strains-For the generation of reporter strains, the upstream regions of the genes *fimA* and *rpoE* were cloned into the pUA66 vector. The restriction sites used are BamHI and XhoI (NEB). The ligated product was transformed into STM WT and STM Δ*malQ*.

### Flow cytometry

Reporter strains were grown in LB medium and subcultured at a 1:33 ratio to obtain a log phase culture. Log phase culture was utilised for FACS. Positive GFP percentage population was analysed using CytoFLEX for 10,000 events.

### Eukaryotic cell lines and growth conditions

Cell line Caco-2 used in this study was maintained in Dulbecco’s Modified Eagle’s Media (Sigma-Aldrich) supplemented with 10% FBS (fetal bovine serum, Gibco), 1% sodium pyruvate, 1% penicillin-streptomycin, and 1% non-essential amino acids at 37°C in the presence of 5% CO_2_. Cells were grown to confluence, and after counting them with a haemocytometer, an equal number of cells were seeded into 24-well plates.

### Invasion assay

One lakh seeded Caco2 cells were infected with bacteria at an MOI of 10, and a log phase culture was used to infect the epithelial cells (Caco2 cells). Cells were centrifuged at 800 rpm for 5 minutes and then incubated at 37°C with 5% CO2 for 25 minutes. Cells were then washed with 1X PBS to remove all extracellular bacteria, and media containing 100µg/ml gentamicin in DMEM with FBS was added for 1 hour. Similarly, 25 µg/ml of gentamicin in DMEM media with FBS was added, and the cells were incubated until lysis. Cells were lysed with 0.1% Triton X-100 at 2 hours. Bacteria were plated on the LB Agar, incubated at 37°C for 12 hours, and colonies were counted. To calculate the percentage invasion, bacterial CFU at 2 hours were divided by the pre-inoculum CFU.

### Adhesion assay

log phase bacteria were adjusted to 0.3 OD and incubated with and without 0.2M Mannose for 1 hour before infection. For adhesion experiments with mannose, the bacteria were kept in static Luria broth at 37°C after subculture to induce type 1 fimbriae (14). Bacteria have been kept at 37°C, 170 rpm for all the other experiments.

Caco2 cells were infected with an MOI of 10 and incubated for 10 minutes. Cells were washed with 1X PBS and lysed using 0.1% Triton X-100. Bacteria were plated on LB Agar and incubated at 37°C for 12 hours, after which the colonies were counted. The number of adhered cells was calculated by dividing the obtained CFU by the pre-inoculum values of the bacteria used for infection. The cells were seeded with coverslips for confocal microscopy, and the assay mentioned above was performed. At the endpoint, the cells were fixed using 3.5% Paraformaldehyde for 10 minutes, followed by washing with 1x PBS and storing the samples in 1x PBS.

### Confocal microscopy

A protocol from Roy Chowdhury et al. was followed (15). The PFA fixed samples from the adhesion assay were stained with a primary antibody against *Salmonella* (1:500 in 2.5% BSA) overnight at 4°C. The primary antibody was washed with 1xPBS, and the secondary antibody was added for 45 minutes at room temperature. Samples were then stained with FM-464 for 10 minutes and subjected to a final wash. Coverslips were mounted on the slides after the addition of mounting media and sealed.

Percentage adhesion was calculated by dividing the number of bacteria by the number of Caco2 cells in the acquired images. Images were acquired using a Zeiss 880 confocal microscope and analysed by Zeiss Zen Black 2012 software.

### Biofilm formation

Bacteria were grown until the stationary phase, inoculated in biofilm media (tryptone and yeast extract) with different concentrations of maltose, and incubated at 28°C for 3 days. The biofilm formed was measured using crystal violet staining, and the absorbance was measured at 570 nm.

### RNA isolation and cDNA Synthesis

STM WT and STM Δ*malQ* were subcultured (1:100) in LB to achieve a log phase for analysis of various invasion and adhesion genes. The bacterial culture was pelleted down by centrifugation at 6000 rpm for 10 min, and 1 ml of TRIzol (Takara) was added to the pellet. Samples were stored at -80°C until they were processed. The lysed supernatant was extracted with chloroform, and RNA precipitation was done using an equal volume of isopropanol. The obtained pellet was washed with 70% ethanol in DEPC water and air-dried. The RNA was dissolved in DEPC water. The quality and quantity of RNA were checked using Nanodrop and 2% agarose gel electrophoresis. DNase treatment (Thermo Fisher Scientific) was performed on the isolated RNA at 37°C for 1 hour, followed by heat inactivation at 65°C for 15 minutes. DNase-treated RNA was converted to cDNA using the PrimeScript Reagent RT kit (Takara). qRT-PCR was performed using the TB Green RT-qPCR Kit in a real-time PCR system. For the analysis of RT data, 16S rRNA was used as the reference gene to calculate the delta Ct (ΔCt) of the samples. ddct was calculated by comparing two different samples.

### Animal experiments

All the animals were acquired from the central animal facility at the Indian Institute of Science, Bangalore. Five-to six-week-old healthy C57BL/6 male mice were used in the study, and they were maintained in stress free environments before infection for 2-3 days, 5 animals per cage.

Maltose supplementation (100 µL) at various concentrations or Milli Q water as control was administered to the mice by oral gavage 1 hour before infection. The mice were then infected with 10^7^ (100 µL) bacteria of STM WT or STM *ΔmalQ* as mentioned in the figure legends. Mice were euthanised 24 hours post-infection, and ileum sections free from faeces were taken, homogenised and plated on *Salmonella Shigella* agar and normalised to organ weight. Mice were euthanised by overdosage of isoflurane ( 2% ) followed by cervical dislocation. 6 animals were used for this experiment for each condition and total 36 animals were used in this experiment. No data points were excluded during data analysis.

To determine the organ burden in mice, mice were orally gavage with 10^7^(100µL) bacteria (STM WT, STM *ΔmalQ,* STM *ΔmalEFG,* STM *ΔmalQ:malQ),* euthanised, and various organs -Blood, Liver, mesenteric lymph node (MLN), spleen were acquired 5^th^ day post infection. Blood was isolated by performing a heart puncture. Mouse organs were homogenised and plated on *Salmonella Shigella* agar to determine the bacterial burden in various organs and normalised to organ weight. 5 animals were used for this experiment for each bacterial strain and total 20 animals were used in this experiment. The animal experiments were not performed in a blind manner, and authors were aware of the different strains used to infect mice in various cages.

### TEM

Transmission electron microscopy was performed using a previously described protocol (16). Stationery and log phase cultures of STM WT and STM Δ*malQ* were washed twice with 1X PBS and resuspended in 50 µL of 1X PBS. 5 µL of the sample was placed on copper grids. After air drying, samples were negatively stained with 1% uranyl acetate. TEM microscopy was performed using a JEM-100CXII JEOL.

### Statistics

The Statistical test, number of biological (N) and technical replicates (n) used in the experiments are mentioned in the figure legends. For the animal experiments, the Mann-Whitney test was performed. All the data were plotted and analysed using GraphPad Prism 8. A *p*-value of <0.05 was considered statistically significant in the experiments.

## Results

### The absence of maltose metabolism increases invasion and adhesion

Previous reports showed that colonising mice with simple sugars utilised by intestinal microbiota decreases the pathogen burden by competing with the pathogen for that sugar (17). To determine the role of maltose in pathogenesis, C57BL/6 mice were treated with various concentrations of maltose. It was observed that at 1% maltose treatment, STM WT infection was reduced in the ileum at 24 hours. This inhibition was not observed at higher maltose concentrations. STM WT and deletion of the maltose hydrolysing gene (STM Δ*malQ)* had similar organ burden in the ileum when no maltose supplementation was provided (**Figure 1A**). Maltose metabolism has been reported to reduce bacterial colonisation from the third day post-infection(7). It was observed that the STM Δ*malQ* showed colonisation defects in blood, mesenteric lymph node (MLN), liver and spleen compared to STM WT at 5 days post-infection (Supplementary Figure 1). However, maltose transporter deletion (STM Δ*malEFG)* showed no growth defect in colonising various mouse organs.

**Figure 1.**
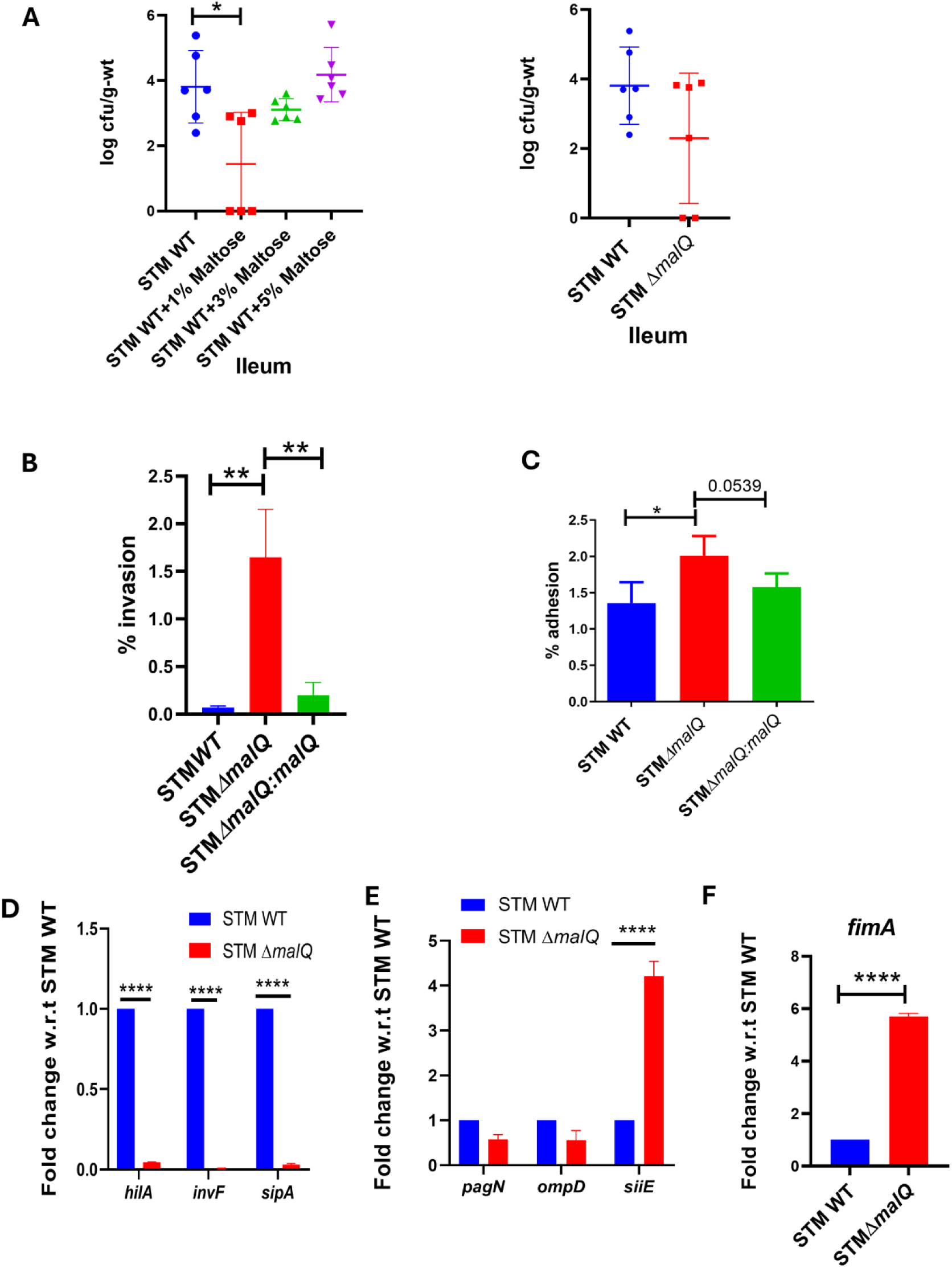
*malQ* deletion increases *Salmonella* invasion and adhesion. A. Organ burden in the ileum of mice upon supplmentation with various concentrations of maltose. The Mann-Whitney test was used for analysis, with six animals per cohort, presented as mean ± SD. B. Percentage invasion of STM WT, STM Δ*malQ,* and STM Δ*malQ:malQ* in Caco2 cells. An unpaired Student’s t-test was used for the analysis. Data are representative of N=3, n=3, presented as mean ±SD. C. Percentage adhesion of STM WT, STM Δ*malQ, and* STM Δ*malQ:malQ* in Caco2 cells. An unpaired Student’s t-test was used for the analysis. Data are representative of N=3, n=3, presented as mean ±SD. D. mRNA expression of SPI-1 genes (*hilA, invF, sipA)* by q-RT PCR, Analysis by two-way ANOVA, Data are representative of N=3, n=3, presented as mean ±SD. E. mRNA expression of large proteins aiding in adhesion (*pagN, ompD, siiE)* by q-RT PCR. Analysis by two-way ANOVA. Data are representative of N=3, n=3, presented as mean ±SD. F. mRNA expression of type 1 fimbriae (*fimA)* by q-RT PCR. An Unpaired Student’s t-test was used for the analysis. Data are representative of N=3, n=3, presented as mean ±SD.

To further study the role of maltose in *Salmonella* infection, upon infection of Caco-2 cells, we observed an increased invasion in Caco-2 cells of STM Δ*malQ,* which is reduced upon complementation of the *malQ* gene (**Figure 1B**). Adhesion to host cells is the first step in facilitating the invasion and colonisation of the host. *Salmonella* adhesion depends on various fimbriae and adhesins secreted by the type 1( BapA and SiiE) and type 5 (MisL, ShdA and SadA) secretion systems. Large outer membrane proteins, such as PagN and Rck, also contribute to adhesion (18). We further examined the adhesion of the *malQ* deleted strain and observed an increased adhesion of Caco2 cells (**Figure 1C**). Upon investigating the mRNA expression of various factors responsible for the invasion and adhesion in the *malQ* deleted strain compared to the STM WT by qRT-PCR. It was observed that SPI-1 genes (*hilA*, i*nvF, sipA),* responsible for the invasion, were downregulated in STM Δ*malQ* compared to STM WT (**Figure 1D**). In the literature, it has been demonstrated that invasion of Caco-2 cells is largely independent of SPI-1, PagN and RcK. The triple knockout of these genes could invade only slightly less than the WT (19). Next, upon investigation of the adhesion genes, encoding large proteins *pagN* and *ompD* were downregulated, whereas *siiE* was upregulated in STM Δ*malQ (***Figure 1E**). *siiE* has been reported to play a role in adhesion only in polarised Caco-2 cells (20). *fimA,* a major structural protein of the type 1 fimbriae that mediates adhesion, was upregulated in STM Δ*malQ* compared to STM WT (**Figure 1F**).

### Deletion of the *malQ* gene increases the expression of type 1 fimbriae

Upon cloning the *fimA* promoter upstream of GFP, we observed that promoter activity was significantly increased in STM Δ*malQ,* resulting in a higher percentage of positive population compared to STM WT (**Figure 2A**). Upon visualisation of type 1 fimbriae with transmission electron microscopy (TEM), more type 1 fimbriae were observed in STM Δ*malQ* bacteria collected from the overnight culture compared to STM WT. Thus, the deletion of *malQ* increased the type 1 fimbriae production (**Figure 2B**). Thus, we concluded that increased *fimA* might be responsible for the increased adhesion and invasion. Our results are in accordance with the literature, which shows an inverse relation between the expression of SPI-1 and type 1 fimbriae (21). The above results demonstrate that the deletion of *malQ* is associated with concomitant increases in fimbriae production, suggesting the anti-adhesive role of *malQ*.

**Figure 2.**
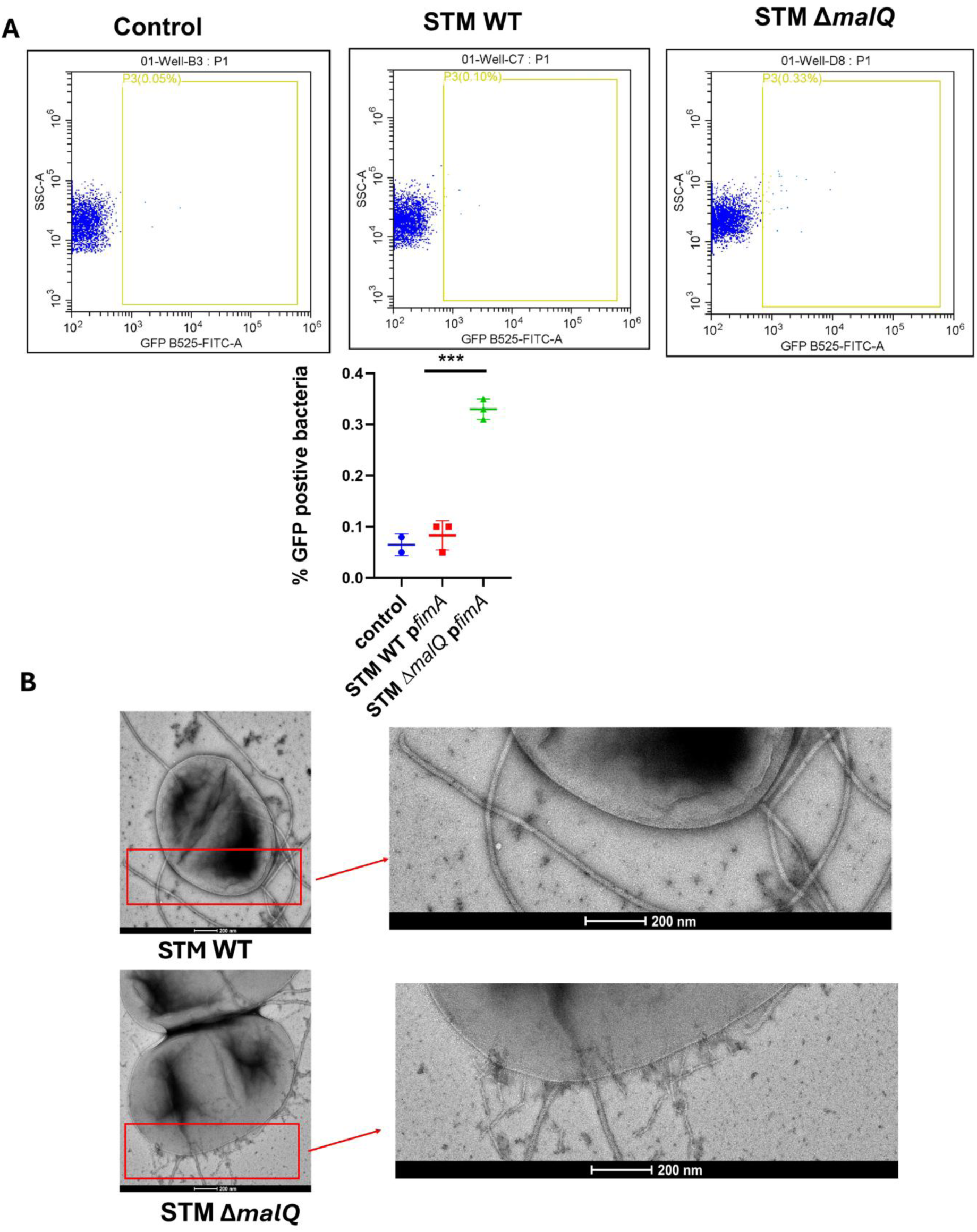
*malQ* deletion in *Salmonella* increases type 1 fimbriae. A. Activity of *fimA* promoter in STM WT and STM Δ*malQ* in the log phase using FACS. Analysis by one-way ANOVA. Data are representative of N=3, n=3, presented as mean ±SD. B. Transmission electron microscopy images of STM WT and STM Δ*malQ*.

### Increased adhesion is due to the enhancement of type 1 fimbriae

Type 1 fimbriae bind to host glycoproteins such as mannose using FimH, and incubation with 0.2 M Mannose can inhibit the binding of *Salmonella* to the host by Type 1 fimbriae (22). To elucidate whether the increased adhesion depended on the interaction with mannose residues of the eukaryotic cells, bacteria were incubated for 1 hour before infection with 0.2M mannose. The increased adhesion of the STM Δ*malQ* was significantly reduced by pretreatment with mannose (**Figure 3A**). Confocal microscopy was performed to confirm further that the increased adhesion in STM Δ*malQ* was due to increased fimbriae. Increased adhesion to Caco2 of STM Δ*malQ* compared to STM WT was observed, which was inhibited by adding mannose (**Figure 3B**). Type 1 fimbriae have also been reported to facilitate biofilm formation (23). The biofilm was grown with various concentrations to determine the role of maltose in biofilm formation. Maltose also acted as an inhibitory signal in biofilm formation. (**Supplementary** Figure 2A). Furthermore, in the biofilm assay with STM Δ*malQ,* STM Δ*malEFG* and STM Δ*lamB,* it was observed that there was no significant reduction of biofilm in STM Δ*malQ* and STM Δ*malEFG,* compared to STM WT (**Supplementary** Figure 2B). Thus, it suggested that the maltose hydrolysis gene *malQ* and maltose transport into bacteria by *malEFG* mediated the decreased biofilm.

**Figure 3.**
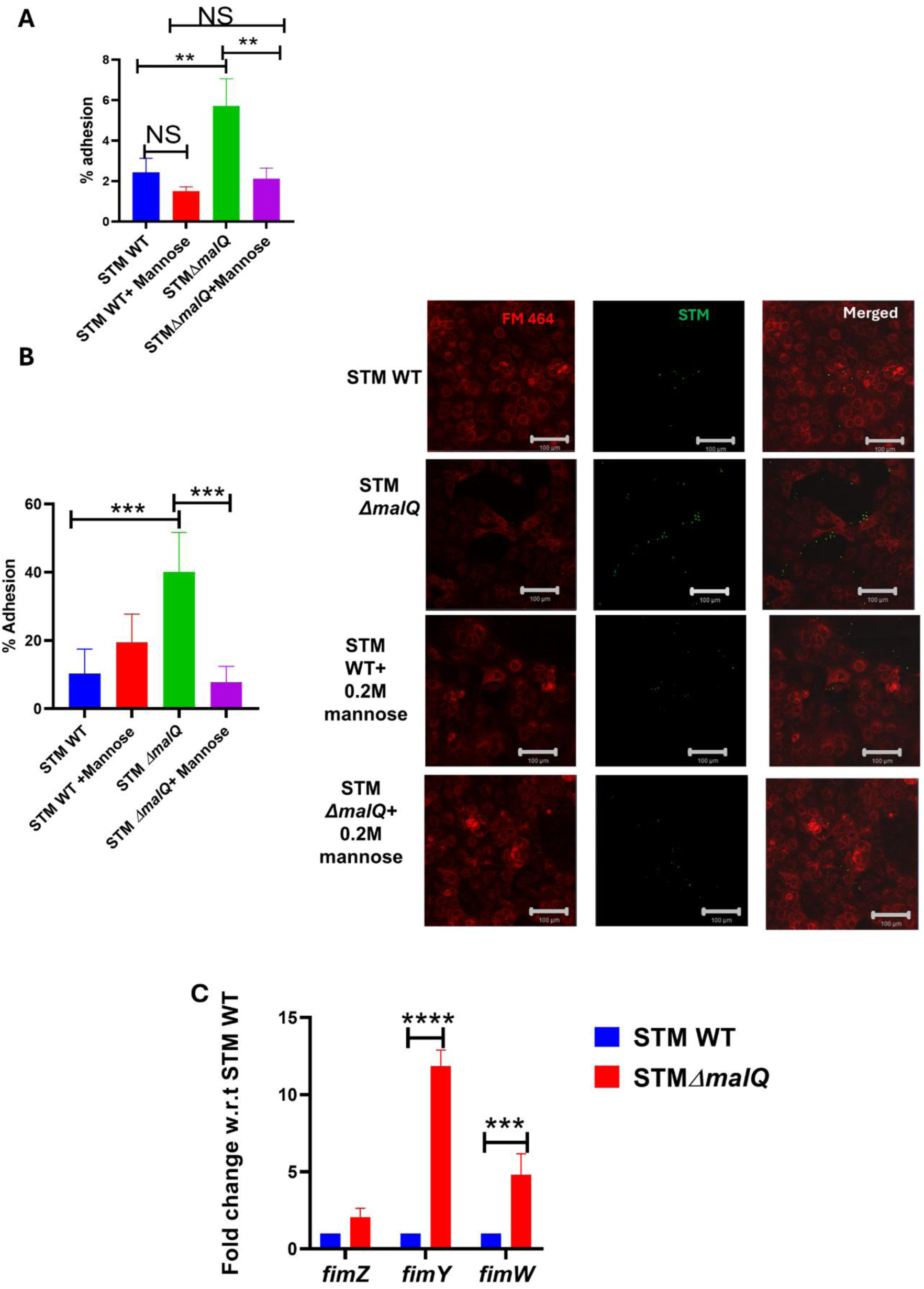
Type 1 fimbriae mediate the increased adhesion of STM Δ*malQ*. A. The percentage adhesion of STM WT and STM Δ*malQ* in Caco-2 cells treated with and without 0.2 M mannose. One-way ANOVA was used for analysis. Data are representative of N=3, n=3, presented as mean ±SD. B. Confocal microscopy was used to determine the adhesion percentage of STM WT and STM Δ*malQ* in Caco-2 cells on treatment with and without 0.2 M mannose. One-way ANOVA was used for the analysis. Data are representative of N=2, n=5, presented as mean ±SD. C. mRNA expression of the fimbrial regulatory molecules *fimZ, fimY* and *fimW* in STM WT and STM Δ*malQ* by using q-RT PCR. Two-way ANOVA was used for the analysis. Data are representative of N=3, n=3, presented as mean ±SD.

To elucidate the mechanism of STM *ΔmalQ*-mediated regulation of type 1 fimbriae. The mRNA expression levels of type 1 fimbriae regulators (*fimY, fimZ* and *fimW*) were determined. FimY and FimZ positively regulate type 1 fimbriae in *Salmonella*, whereas FimW negatively regulates them. FimZ also represses the expression of SPI-1 via *hilE* (24, 25). There was no change in the expression of *fimZ,* whereas *fimY* and *fimW* were significantly upregulated in the STM *ΔmalQ* (**Figure 3C**). The significantly increased expression of *fimY* might be responsible for the increased production of type 1 fimbriae.

### RpoE regulates type 1 fimbriae

Type 1 fimbriae in Gram-negative bacteria, such as uropathogenic *E.coli* and *Serratia marcescens,* are reported to be repressed by the carbohydrate metabolism-regulating system cAMP-CRP (26, 27). Upon analysis of the mRNA expression of CRP in STM *ΔmalQ,* we observed significant downregulation compared to STM WT (**Figure 4A**). Thus, the increased type 1 fimbria production in STM *ΔmalQ* was due to increased expression of *fimY,* which might be modulated by the decreased expression of CRP. Previous reports have shown that the *rpoE* promoter 1 may be negatively regulated by cAMP-CRP in *Salmonella,* inducing promoter 3 of rpoE during carbon starvation (28). *rpoE* also regulates the pathogenicity genes of *E.coli* in interaction with human brain microvascular endothelial cells and the thin aggregative fimbriae(Agf) in *Salmonella* (29, 30). The qRT-PCR analysis of the *rpoE* gene in STM Δ*malQ* revealed upregulation of *rpoE* compared to STM WT (**Figure 4B**). Similarly, for the promoter activity of *rpoE*, we cloned promoter 3 of *rpoE* upstream of GFP and observed that STM Δ*malQ* has a significantly higher percentage positive population expressing *rpoE* compared to STM WT (**Figure 4C**). Thus, *malQ* deletion leads to increased promoter activity and transcript levels of *rpoE*. To determine the role of *rpoE* in the increased adhesion of STM *ΔmalQ,* we deleted *rpoE* in STM Δ*malQ* and observed that the increased adhesion of STM Δ*malQ* was lost when performing adhesion to Caco2 cells. STM Δ*rpoE* had less adhesion than STM WT, and double deletion of *rpoE* and *malQ* resulted in adhesion similar to that of STM Δ*rpoE* (**Figure 4D**). Thus, the increased adhesion of STM *ΔmalQ* was mediated by the alternative sigma factor RpoE.

**Figure 4.**
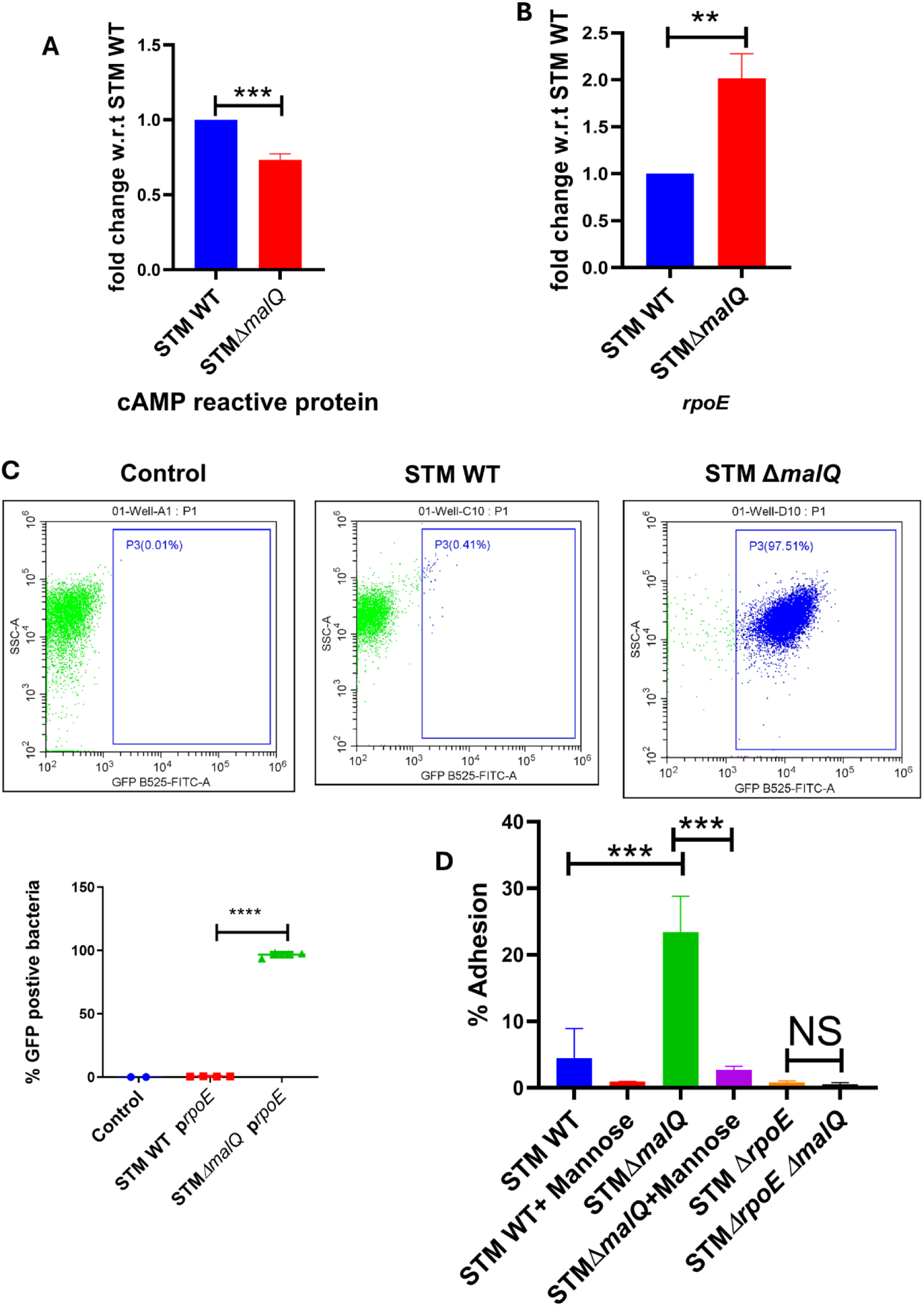
Role of *rpoE* in type 1 fimbriae regulation. A. mRNA expression of cAMP reactive protein (CRP) in STM WT and STM Δ*malQ* using q-RT PCR. An unpaired Student’s t-test was used for analysis. Data are representative of N=3, n=3, presented as mean ±SD. B. mRNA expression of *rpoE* in STM WT and STM Δ*malQ* using q-RT PCR. An unpaired Student’s t-test was used for analysis. Data are representative of N=3, n=3, presented as mean ±SD. C. Activity of Promoter3 of *rpoE* in STM WT and STM Δ*malQ* in the log phase using FACS. Analysis by one-way ANOVA. Data are representative of N=3, n=4, presented as mean ±SD. D. Percentage adhesion of STM WT, STM Δ*malQ* in Caco-2 cells on treatment with and without 0.2 M mannose, STM Δ*rpoE* and STM Δ*rpoE* Δ*malQ.* One-way ANOVA was used for the analysis. Data are representative of N=3, n=3, presented as mean ±SD.

## Discussion

Dietary polysaccharides in food are broken down into disaccharides (maltose, sucrose, and lactose). *Salmonella* is generally considered unable to utilise disaccharides such as sucrose and lactose; however, serovars like *Salmonella* Mbandaka and *Salmonella* Bareilly can ferment sucrose (31, 32). Recent reports have also shown that certain *Salmonella* Typhimurium types can utilise sucrose due to the acquisition of genes important for metabolism (31). Antibiotic-resistant lactose fermenters of *Salmonella* Typhimurium have been reported in cattle (32). Compared to sucrose and lactose, the most utilised disaccharide sugar, maltose, in the intestine by *Salmonella,* has been studied less in its role in pathogenesis. In this study, we investigated the role of maltose in modulating the virulence of *Salmonella*.

Maltose utilisation is essential for the survival of *E.coli* in the mouse intestine, as shown by Jones et al. (7). As observed earlier, we found no difference in the intestinal colonisation of maltose metabolism mutant STM Δ*malQ* in the mouse’s ileum after 1-day post-infection. However, decreased ileum colonisation of STM WT was observed in mice treated with 1% maltose. Furthermore, STM Δ*malQ* showed higher invasion and adhesion in the epithelial cells Caco-2. The higher adhesion depended on the higher expression of type 1 fimbriae by STM Δ*malQ.* The Inhibition of adhesion mediated by type 1 fimbriae with the addition of mannose reversed the advantage of increased adhesion. The mechanism shown by Muller et al. involves the Carbohydrate metabolism-regulating system cAMP-CRP, which mediates type 1 fimbriae expression through fimB-mediated recombination in *E.coli,* a process absent in *Salmonella* (*26*). Further studies are required to elucidate how the maltose metabolism gene (*malQ*) regulates fimbriae production through the cAMP-CRP pathway. We attempted to overexpress CRP in STM Δ*malQ* but were unable to do so, as the induction was toxic to the bacteria. The alternative sigma factor *rpoE* is essential for maintaining the membrane integrity of bacteria and for coping with various stresses they face (33). RpoE has also been known to induce fimbriae and invasion in *E.coli* (29). We observed higher mRNA expression and promoter activity of *rpoE* in STM Δ*malQ* compared to STM WT. Deletion of the *rpoE* in STM Δ*malQ* had a similar effect to the addition of mannose, and STM Δ*rpoE,* STMΔ*malQ*Δ*rpoE* had similar adhesion. Thus, the increased adhesion of STM Δ*malQ* is dependent on the presence of *rpoE*.

In conclusion, we have demonstrated that the maltose-hydrolysing gene (*malQ*) reduces invasion and adhesion to Caco-2 epithelial cells. The reduced adhesion is mediated by the reduced expression of type 1 fimbriae, which is mediated by RpoE. Therefore, our study reveals a novel link between dietary sugar and the adhesion of the foodborne pathogen *Salmonella*.

## Funding

This work was funded by the DAE SRC fellowship and the DBT-IISc partnership program for advanced research in biological sciences and Bioengineering to DC. We acknowledge the infrastructure support from the ICMR (Centre for Advanced Study in Molecular Medicine), DST (FIST), TATA Fellowship, and UGC (Special Assistance). KP sincerely acknowledges IISc-MHRD for the fellowship. The funders played no role in the study design, data collection, analysis, or manuscript writing.

## Ethics statement

All animal experiments were approved by the Institutional Animal Ethics Committee, Indian Institute of Science, Bangalore, India. The ethical clearance number for this study was CAF/ethics/852/2021. All experiments were conducted in strict accordance with the guidelines provided by National Animal Care and the Committee for the Purpose of Control and Supervision of Experiments on Animals (CPCSEA, a statutory Committee, established under Chapter 4, Section 15 (1) of the Prevention of Cruelty to Animals Act 1960). (CPCSEA Registration No. 435 48/1999).

## CRediT authorship contribution statement

**Kirti Parmar**-writing-review and editing, visualisation, experimental design, project administration, investigation and data analysis. **Jayashree S-** Investigation. **Dipshikha Chakravortty**-writing-review and editing, visualisation, experimental design, investigation, data analysis, resources and supervision.

## Acknowledgements

Abhilash Vijay Nair is acknowledged for helping with the oral gavage of the animals. We acknowledge the department’s Real-time facility, confocal microscopy facility and central animal facility at IISc for supporting the experiments.

## Declaration of competing interest

The authors declare no conflict of interest.

## Data and material availability

All the data are available in the main text or the supplementary file.

## Supplementary information

**Supplementary Figure S1.**
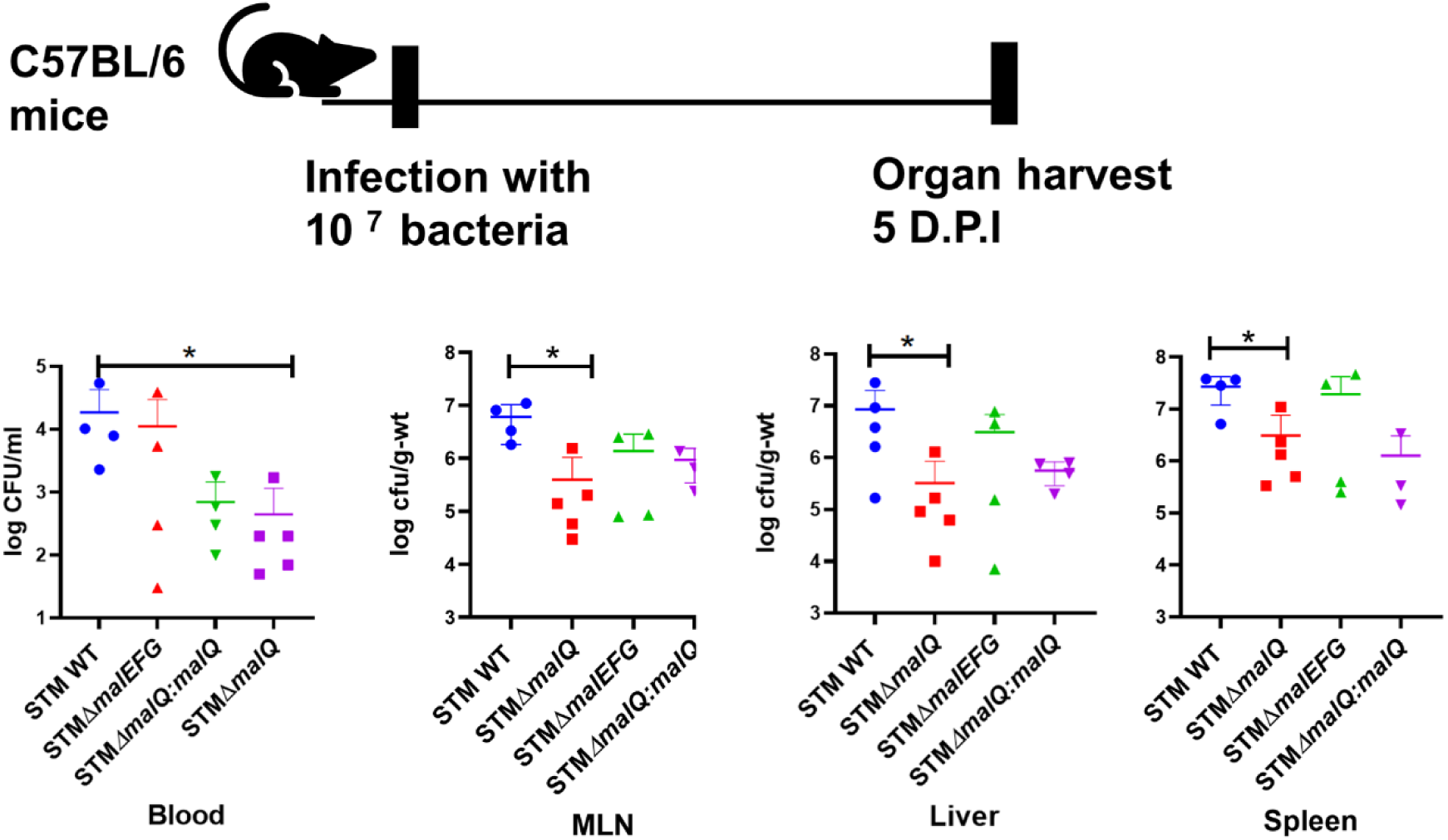
Maltose metabolism gene *malQ* aids in the colonisation of mice. *In-vivo* pathogenesis of STM WT, STM Δ*malQ* and STM Δ*malEFG* in C57BL/6 mice 5 days post-infection. Organ burden in blood, MLN, Liver and spleen was determined. The Mann-Whitney test was performed for the analysis, with five animals per cohort, and the results are presented as mean ± SD.

**Supplementary Figure S2.**
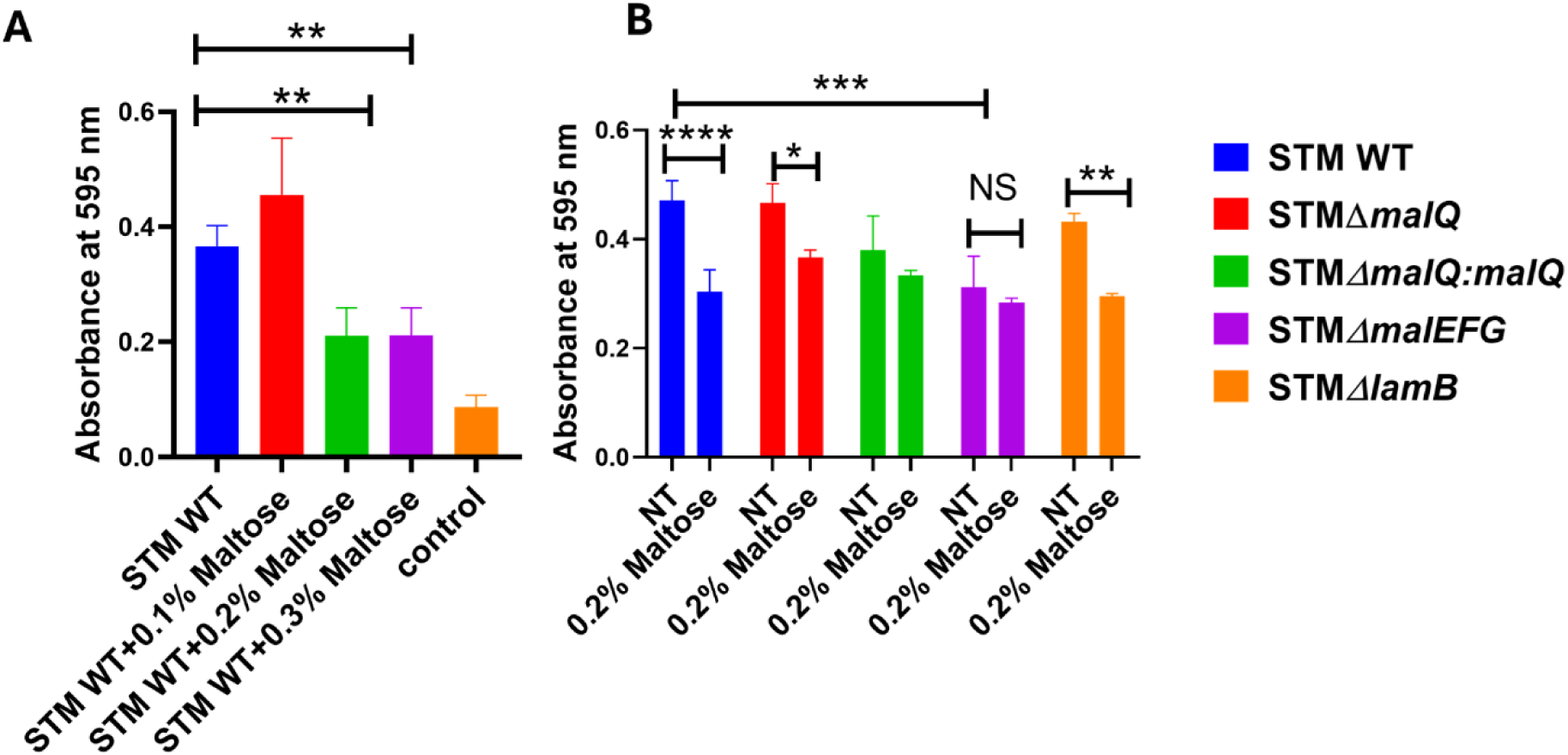
Maltose inhibits the biofilm formation of *Salmonella* Typhimurium. A. Crystal violet staining of the biofilm 3 days post inoculation with different concentrations of maltose. Data is representative of N=3,n=4, presented as mean ± SD, analysis by unpaired t-test. B. Crystal violet staining of the biofilm 3 days post inoculation with STM WT, STM STM Δ*malQ,* STMΔ*malEFG* and STM Δ*lamB* on treatment with and without treatment of 0.2 % maltose. The data are representative of N = 3, n = 4, and are presented as mean ± SD. Analysis was performed using two-way ANOVA.

**Supplementary Table S1.** List of bacterial strains used in the study.

| S.NO | Bacterial strain | Antibiotic |
| --- | --- | --- |
| 1 | STM WT 14028S | - |
| 2 | STM $\Delta$ *malQ* | Kanamycin/<br>Chloramphenicol |
| 3 | STM $\Delta$ *malEFG* | Kanamycin |
| 4 | STM $\Delta$ *lamB* | Chloramphenicol |
| 5 | STM $\Delta$ *malQ*: *malQ* | Kanamycin:<br>Chloramphenicol |
| 6 | STM $\Delta$ *rpoE* $\Delta$ *malQ* | Kanamycin:<br>Chloramphenicol |
| 7 | STM $\Delta$ *rpoE* (Made for earlier study) (34) | Chloramphenicol |
| 8 | STM WT *PrpoE*-GFP | Kanamycin |
| 9 | STM $\Delta$ *malQ* *PrpoE*-GFP | Kanamycin:<br>Chloramphenicol |
| 10 | STM WT *PfimA*-GFP | Kanamycin |
| 11 | STM $\Delta$ *malQ* *PfimA*-GFP | Kanamycin:<br>Chloramphenicol |

**Supplementary Table S2.**
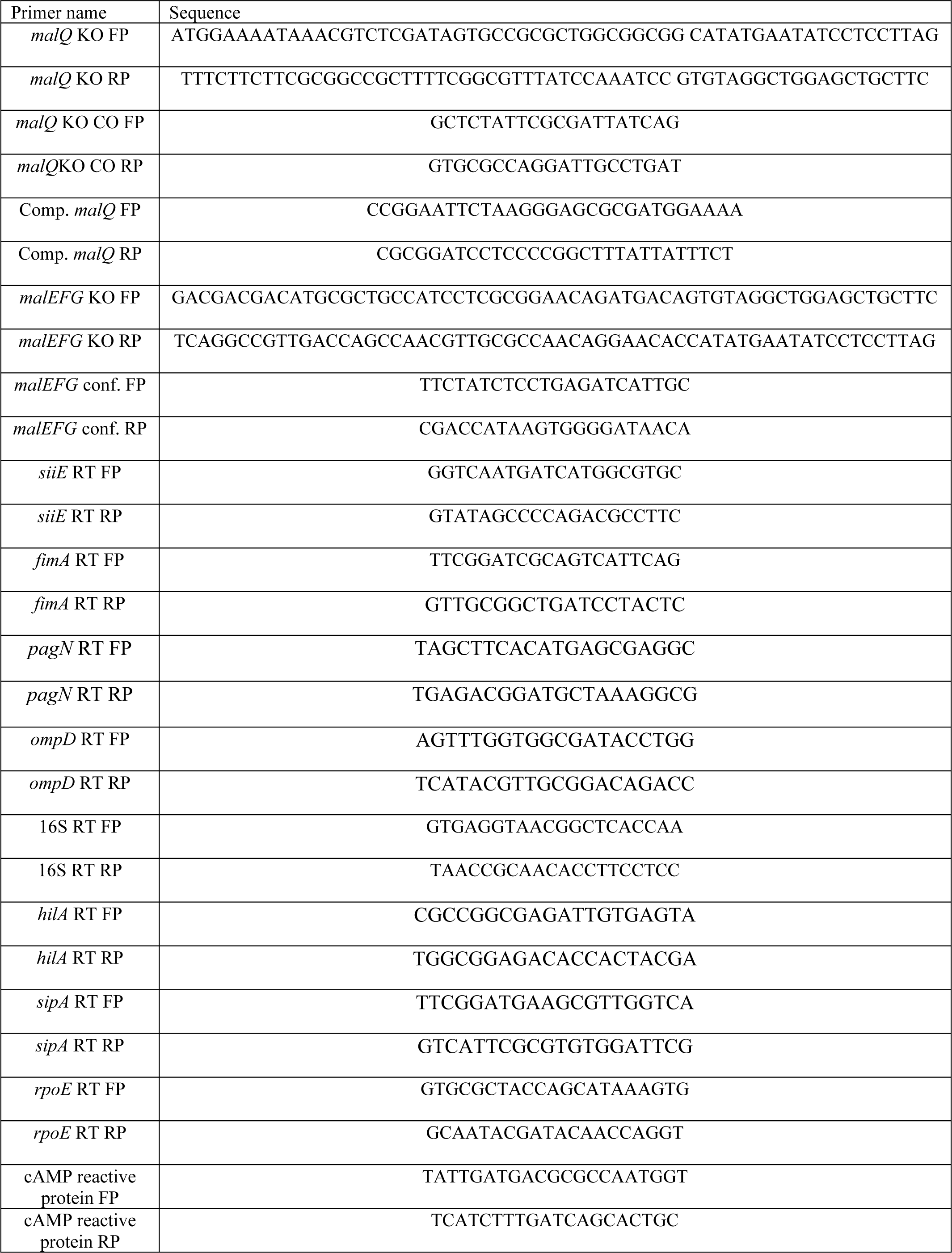
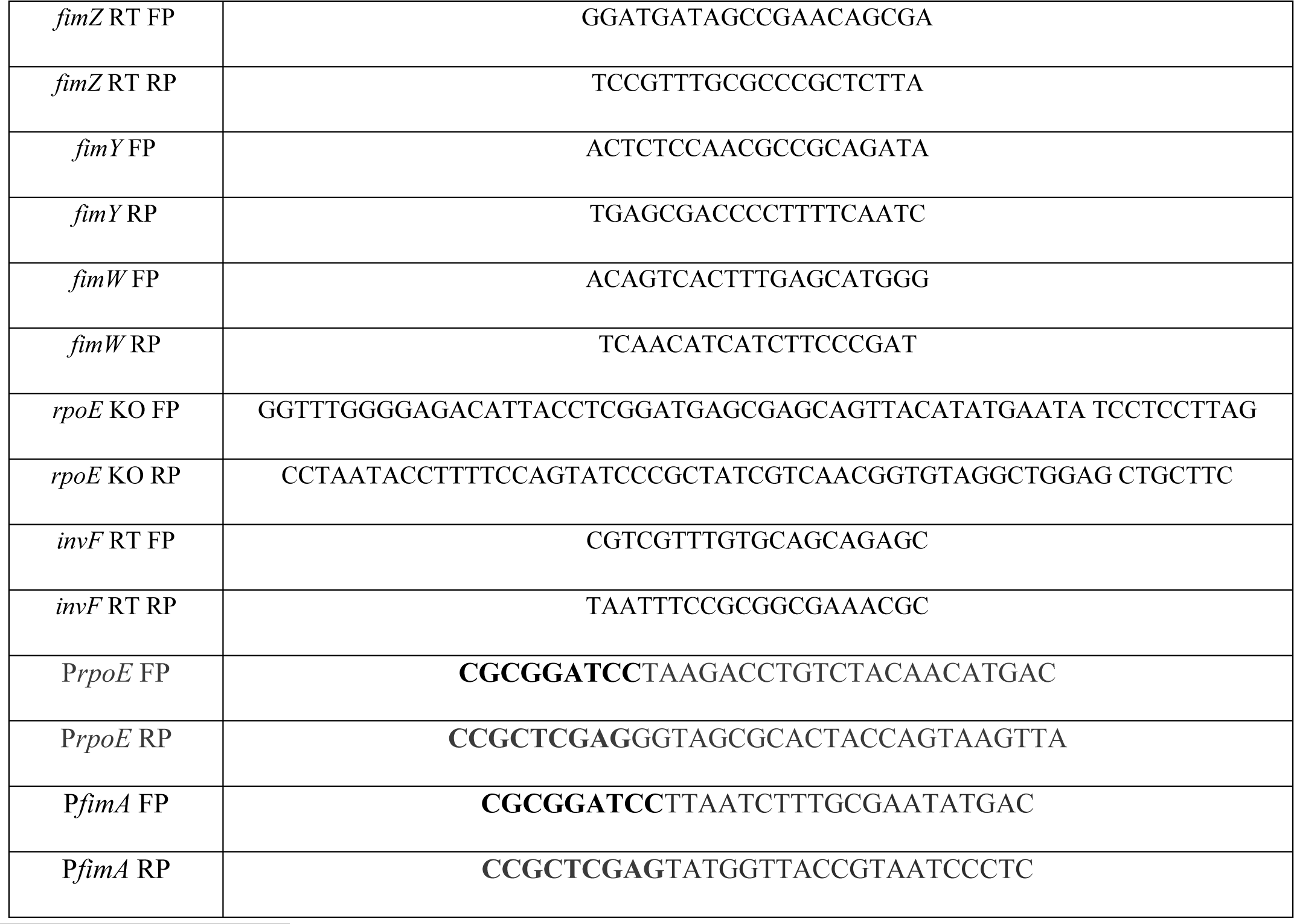
List of primers used in the study.

## Notes

### Competing Interest Statement

The authors have declared no competing interest.

### Summary of Updates

We have revised the manuscript to separate the original study into two distinct papers. The previous version combined two topics- the role of maltose in inhibiting adhesion to epithelial cells and its metabolic advantage under nitrogen-deficient conditions. This current manuscript now concentrates on the inhibitory effect of maltose on epithelial cell adhesion and has been updated with new experimental results to strengthen this focus. The section regarding the advantage of maltose in nitrogen-poor environments has been removed and will be thoroughly investigated in a separate, forthcoming manuscript.

